# Empirical optimization of dual-sgRNA design for in vivo CRISPR/Cas9-mediated exon deletion in mice

**DOI:** 10.1101/2025.09.08.675005

**Authors:** Sung-Yeon Lee, Seongwon Ma, Sangjun Davie Jeon, Hyoju Kim, Beomjoon Jo, Seung-Hoon Han, Eunsoo Jang, Jimin Lee, Yong-Kyu Lee, Dasom Lee

## Abstract

CRISPR/Cas9 has transformed gene editing, enabling precise genetic modifications across species. However, existing sgRNA design prediction models based on in vitro data are difficult to generalize to in vivo contexts. In particular, approaches based on single-sgRNA design require additional filtering of in-frame mutations, which is inefficient in terms of both time and cost. In this study, we developed the first mammalian in vivo-trained prediction model to evaluate the efficiency of a dual-sgRNA-based exon deletion strategy. Using 230 editing outcomes of postnatal viable individuals, eight prediction models were constructed and evaluated based on generalized linear models and random forests. The final selected model, a Combined GLM, integrated the DeepSpCas9 score with k-mer sequence features, achieving an AUC of 0.759 (95% Confidence Interval: 0.697–0.821). Motif analysis revealed that CC sequences were associated with high efficiency and TT sequences were associated with low editing efficiency. This study demonstrates that integrating sequence-based features with existing design scores can improve sgRNA efficiency prediction in vivo. The proposed framework can be applied to the development of next-generation sgRNA design tools, with direct implications for gene therapy, effective animal model generation, and precision genome engineering.

**GRAPHICAL ABSTRACT:** 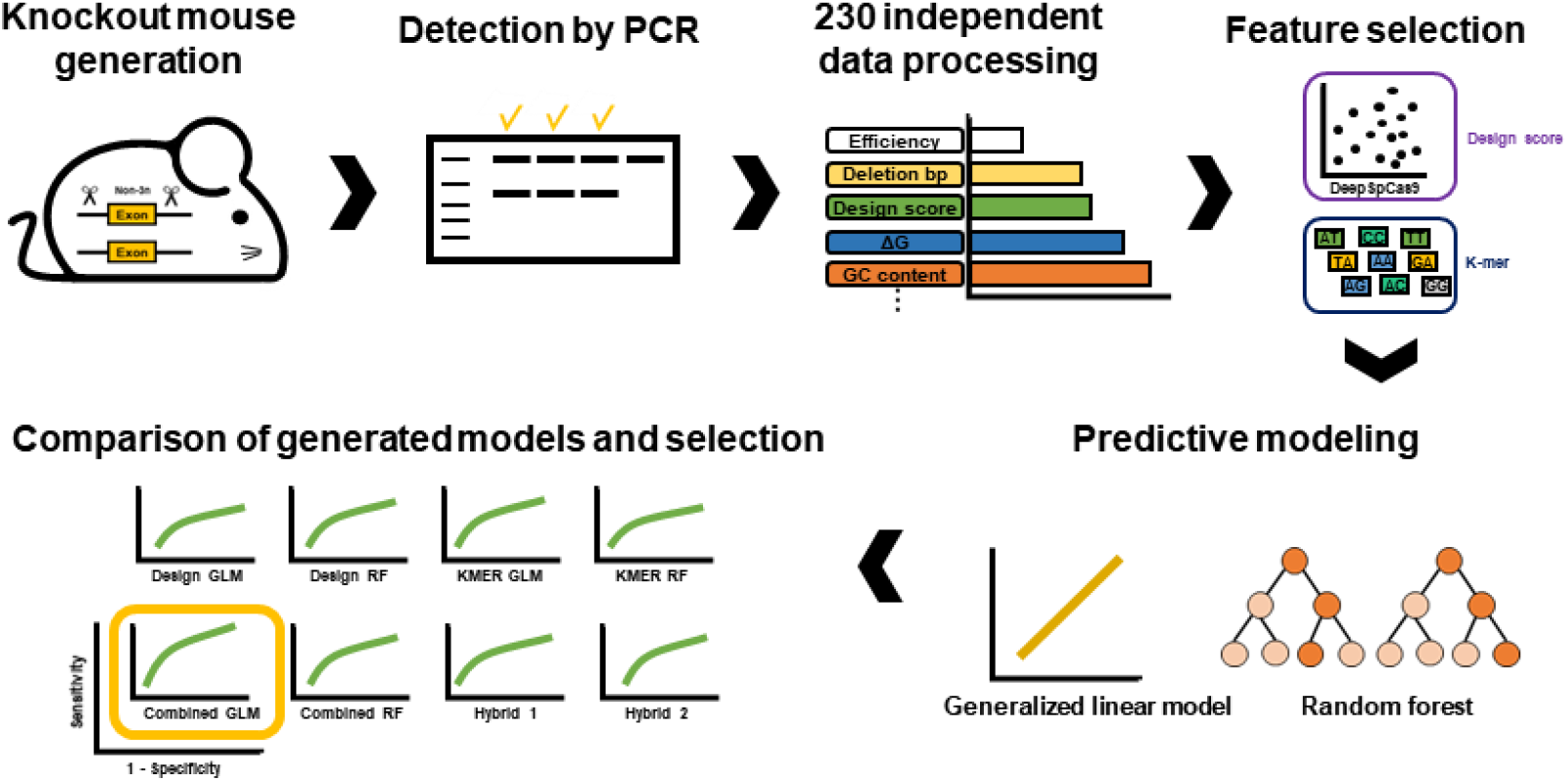

## INTRODUCTION

CRISPR technology has been widely applied across diverse gene editing fields [1–5]. More precise technology such as base and prime editing has been developed to enhance accuracy and reduce off target effects [6, 7], with traditional approaches involving double-strand breaks remaining widely used for their simplicity and robust efficiency [8–12].

Efficient delivery of CRISPR components into embryos is also crucial for successful gene editing. These components are typically introduced via plasmid-based systems or ribonucleoprotein (RNP) complexes, with RNPs preferred for their lower risk of genomic integration and reduced off-target activity [3, 13, 14].

Consequently, various sgRNA efficiency prediction tools have been introduced. Most are trained in vitro and focus on single-sgRNAs [15–22]. However, with respect to achieving gene knockout, these approaches often yield small indels, and in-frame mutations (multiples of 3 bp) may retain gene function, necessitating additional validation methods such as molecular cloning or NGS [23, 24], which increase experimental cost and complexity.

To address these challenges, dual-sgRNAs targeting non-3n exons can promote exon deletion or skipping, ensuring functional gene knockout. This strategy enables straightforward validation via PCR and Sanger sequencing, and can be evaluated in silico beforehand, incorporating validated transcripts in CRISPR-induced indel distributions [25–27]. Such features are particularly advantageous in large animal models or clinical contexts, where ethical and financial constraints demand high precision.

Nevertheless, systematic in vivo datasets supporting dual-sgRNA-mediated knockout efficiency remain limited, and existing models lack training in mammalian in vivo contexts, restricting their predictive power.

Herein, we present the first in vivo-trained prediction model for dual-sgRNA-mediated gene knockout in mammals. Based on five existing prediction tools and k-mers, eight predictive models employing generalized linear models (GLMs) and random forests (RFs) were developed and evaluated. The final model outperformed conventional single-sgRNA-based methods in both prediction accuracy and sgRNA selection. This tool enables accurate in silico prediction of editing outcomes, providing practical guidance for genome editing experiments and supporting more efficient, cost-effective knockout strategies.

## MATERIALS AND METHODS

### Ethics

Due to confidentiality agreements, the nucleotide sequences used for modeling cannot be disclosed. All experiments were conducted at GEM division of Macrogen Inc. (Seoul, Republic of Korea), in accordance with the guidelines of the Korean Food and Drug Administration. The protocols underwent review and approval by the Institutional Animal Care and Use Committees (IACUC; MS-2018-01, MS 2019-01, MS-2020-01, MS-2021-01, MS-2022-01, MS-2023-01, and MS-2024-01). All experiments were carried out in adherence to applicable legal guidelines.

### Design of dual sgRNAs

All designs were based on transcript and FASTA sequences retrieved from the NCBI and Ensembl databases. sgRNAs were designed within intronic regions flanking non-3n exons to induce gene knockout, with exon skipping as the primary objective. To avoid unintentional skipping of adjacent exons located beyond the targeted introns, splicing regulatory elements were carefully considered: splicing donors in the 5′ intron and splicing branch points and acceptors in the 3′ intron. Off-target analysis was conducted using CHOPCHOP v3.0.0 (https://chopchop.cbu.uib.no) [21], CRISPOR v5.2 (https://crispor.gi.ucsc.edu) [22], and the RGEN (http://www.rgenome.net/mich-calculator/) [20] tool, with a preference for sgRNAs showing zero predicted off-targets with up to three mismatches. In cases where this was not feasible, sgRNAs with off-targets located in intronic or intergenic regions were selected. To facilitate genotyping, regions containing mononucleotide or dinucleotide repeats, as well as regions with high AT or GC content surrounding the sgRNA target sites, were excluded. The strand orientation of the sgRNA was not considered.

### Knockout mouse generation

C57BL/6 and ICR mice purchased from Orientbio Inc. (Sungnam, Republic of Korea) were housed at GEM center of Macrogen Inc. in a specific pathogen-free environment. Mice were kept in controlled environment with temperature and humidity maintained at 22 ± 1 °C and 50%, respectively, in 12-hour light and dark cycle. Mice were given free access to feed and water and housed in sterilized individually ventilated cages.

Target sgRNA were synthesized using GeneArt™ Precision gRNA Synthesis Kit (Thermo Fisher Scientific, USA) according to the manufacturer’s instructions. For superovulation, PMSG (Pregnant mare serum gonadotropin; Dsmbio, Republic of Korea) and hCG (Human chorionic gonadotropin; Dsmbio, Republic of Korea) were treated in C57BL/6 female mice. PMSG (7.5IU) was intraperitoneally administered followed by hCG (7.5IU) injection after 48 hours in female mice of 5-8 weeks of age. After hCG injection, these female mice were mated with C57BL/6 stud male mice. 20 hours later, females visually confirmed for copulation plug were sacrificed, and zygotes were collected from the oviduct. Zygotes were then washed in Quinn’s Advantage™ Medium with HEPES (Cooper surgical, Denmark) medium and transferred to KSOM medium (Sigma-Aldrich, USA) droplets in CO_2_ incubation at 37°C. Electroporation was performed 24 hours after hCG treatment at 1 cell stage.

Electroporation was performed using NEPA 21 electroporator (Nepa Gene, Ichikawa, Japan) with RNP complex consisting of SpCas9 protein (Macrogen; 0.62 pmoles/µl) and purified sgRNA pair (each 0.77 pmoles/µl) with Quinn’s Advantage™ Medium with HEPES. The following parameters were used for electroporation: poring pulse: voltage, 225 V; pulse length, 1.5 ms; pulse interval, 50 ms; number of pulses, 4 and transfer pulse: voltage, 20 V; pulse length, 50 ms; pulse interval, 50 ms; number of pulses, 5. Then, embryos were incubated at 37℃ for 2 hours and transplanted by surgical methods into the oviducts of the pseudopregnant recipient mice (ICR).

For founder mice screening, genotyping was performed using DirectPCR Lysis Reagent (102-T; Viagen, USA) according to the manufacturer’s instructions. Then, PCR assay was conducted using Axen™ High-Q Taq DNA Polymerase (MBAE-106; Macrogen) and EF-Taq DNA Polymerase (SEF16-R50h; Solgent, Republic of Korea).

### Data processing

To select sgRNA pair datasets, editing efficiency was first evaluated based on the number of positive pups relative to the total number of pups born, distinguished by PCR-based size differences. For modeling purposes, only autosomal targets were included to avoid allelic imbalance, and data were collected only from experiments in which more than ten pups were obtained to ensure reliability. All experiments were performed in independent batches, and only datasets generated using dual-sgRNAs were included in the analysis.

For each selected sgRNA pair, key sequence-based features were recorded, including microhomology score, GC content, free energy change (ΔG), deletion size, and strand orientation. In addition, cleavage efficiency scores were documented from widely used prediction tools such as RuleSet2 [18], RuleSet3 [19], DeepSpCas9 [16], CRISPRon [15], and CRISPRscan [17]. The final metadata included the number of embryos used, the number of surrogates, the total number of pups born, and the number of offspring confirmed to carry the intended deletion, while out-of-frame indel scores were excluded.

### Correlation, comparative, and distribution analyses

To evaluate the relationship between sgRNA design features and genome editing efficiency, correlation, association, and distribution analyses were performed using a curated dataset collected from dual-sgRNA experiments. For all variables except deletion size (bp) and strand orientation, the minimum value obtained between each sgRNA within the pair was used in the analysis to represent the pairwise design characteristics.

The thermodynamic stability of sgRNA molecules was estimated by calculating the minimum ΔG of each guide RNA using RNAfold, which included both the spacer and scaffold sequences. In addition, to estimate the duplex stability when both sgRNAs are expressed simultaneously, the hybridization free energy (ΔG_cofold) between paired sgRNAs was calculated using RNAcofold.

For correlation analysis, Pearson correlation coefficients were calculated to evaluate linear relationships between editing efficiency and various sequence-based and predictive features. The variables analyzed included microhomology, GC content, ΔG, ΔG cofold, deletion bp, and sgRNA affinity prediction scores from several algorithms, which include RuleSet2, RuleSet3, DeepSpCas9, CRISPRon, and CRISPRscan.

For comparative analysis, the dataset was divided into high and low-efficiency groups based on the median of editing efficiency. The same variables used in the correlation analysis were compared between the two groups using Welch’s t-test, and statistical significance was annotated on boxplots. In addition, sgRNA strand orientation (Convergent, Divergent, and Same) was treated as a categorical variable, and the Wilcoxon rank-sum test was applied to assess whether relative orientation affected editing efficiency.

Finally, to visually examine the overall distribution of each design feature, density plots were generated. A total of ten variables were included in the analysis: five sgRNA prediction scores (RuleSet2, RuleSet3, DeepSpCas9, CRISPRon, CRISPRscan), three sequence-based features (Microhomology, GC content, ΔG), deletion bp, and efficiency.

### K-mer-based profiling and labeling

To characterize nucleotide composition patterns within sgRNA sequences, a k-mer-based profiling approach was employed. Specifically, 2-mers (k = 2) were extracted from target sequences (crRNA) of both sgRNA1 (sg1) and sgRNA2 (sg2) sequences for each data. The extracted k-mers were compiled into a long-format table, corresponding label, and individual k-mer entries. The frequencies of k-mers from the sgRNA pair were computed and reshaped into a wide-format matrix, with columns representing specific 2-mers and rows corresponding to data.

For classification purposes, data were labeled as “High” or “Low” based on whether their editing efficiency was greater than or equal to the median value across all data. The final dataset consisted of frequency profiles of 2-mers along with binary efficiency labels and was used for downstream machine learning analysis.

### Predictive modeling using efficiency-based binary classification

Logistic regression was applied due to its suitability in low-dimensional binary classification problems, where the number of covariates is small relative to the data size [28]. By applying binary classification (High vs. Low) based on the median value of gene editing efficiency, eight GLM- and RF-based models were developed using sgRNA design scores and sequence-based k-mer profiles. The modeling framework consisted of four main stages.

#### Design feature-based modelling

Among the sgRNA design features, variables highly correlated with efficiency were selected, and the minimum value between each sgRNA (sg1 and sg2) was used as input variables. These were then used to train GLM and RF models.

#### K-mer-based modelling

For each independent experiment, all possible 2-mer sequences were extracted from both sg1 and sg2 to generate k-mer frequency matrices, which were then used as input features for training GLM and RF models. Feature importance was assessed using absolute coefficient values for GLM and mean decrease in Gini for RF. To further examine the distribution and significance of k-mers, all k-mers used in the GLM were extracted, and each sgRNA sequence was decomposed into 2-mers with positional information.

#### Combined modeling (Design feature + K-mer)

To enhance prediction accuracy, design features and k-mer profiles were combined into a single feature set. Combined GLM and RF models were trained. Feature importance analyses were performed to identify key variables contributing to predictive performance.

#### Hybrid modelling

To explore potential synergy between modeling strategies, two hybrid models were constructed by averaging prediction probabilities from complementary models. Hybrid 1 combined predictions from the design-based GLM and the k-mer-based RF, while Hybrid 2 combined the design based RF and the k-mer-based GLM.

For each model, Receiver Operating Characteristic (ROC) curve data were constructed by calculating sensitivity and specificity from the predicted values to evaluate predictive performance.

### Model evaluation and selection

To identify the optimal predictive model for genome editing efficiency, we evaluated eight classification models—including GLM and RF variants—based on design features, k-mer features, and their combinations, along with two hybrid models incorporating selected top-performing variables. Model performance was primarily assessed using the AUC, calculated on both hold-out test sets and repeated 10-fold cross-validation.

For further analysis, Pearson’s and Spearman’s correlation analyses were conducted to compare predicted and observed efficiency. Pearson’s R and Spearman’s R were computed using predictions from repeated 10-fold cross-validation to evaluate, respectively, linear and monotonic rank-based relationships with the true outcomes. Additional evaluation metrics included accuracy, sensitivity, specificity, F1 score, balanced accuracy, calibration curves, Brier scores, and precision–recall (PR) curves.

### Statistics

All statistical analyses were conducted in R (v4.5.1). Pearson correlation coefficients were used to assess linear relationships between continuous variables. Welch’s t-test was applied to compare editing efficiency between groups, and the Wilcoxon rank-sum test was used to assess differences in strand orientation. ROC-based AUC values were compared between models using Welch’s t-test and DeLong’s test. Calibration performance was assessed using Brier score and Hosmer–Lemeshow goodness-of-fit test, along with calibration and PR curves to evaluate prediction reliability. For each k mer, Fisher’s exact test was used to compare its frequency between high- and low-efficiency groups, followed by multiple testing correction using Benjamini–Hochberg method. K-mers with False Discovery Rate (FDR) below 0.05 were considered statistically significant.

## RESULTS

### Data collection and basic statistical analysis

In total, 230 independent in vivo experimental outcomes were obtained, comprising 192 types of genes and 201 sgRNA pairs. The overall workflow of our study is visualized in **Figure 1A**. On average, 71.9 embryos were used per editing experiment, with approximately 3.31 surrogate recipients used per case. Each experiment yielded an average of 23.45 offsprings, among which 5.84 were confirmed to carry the intended deletion (**Figure 1B**).

**Figure 1.**
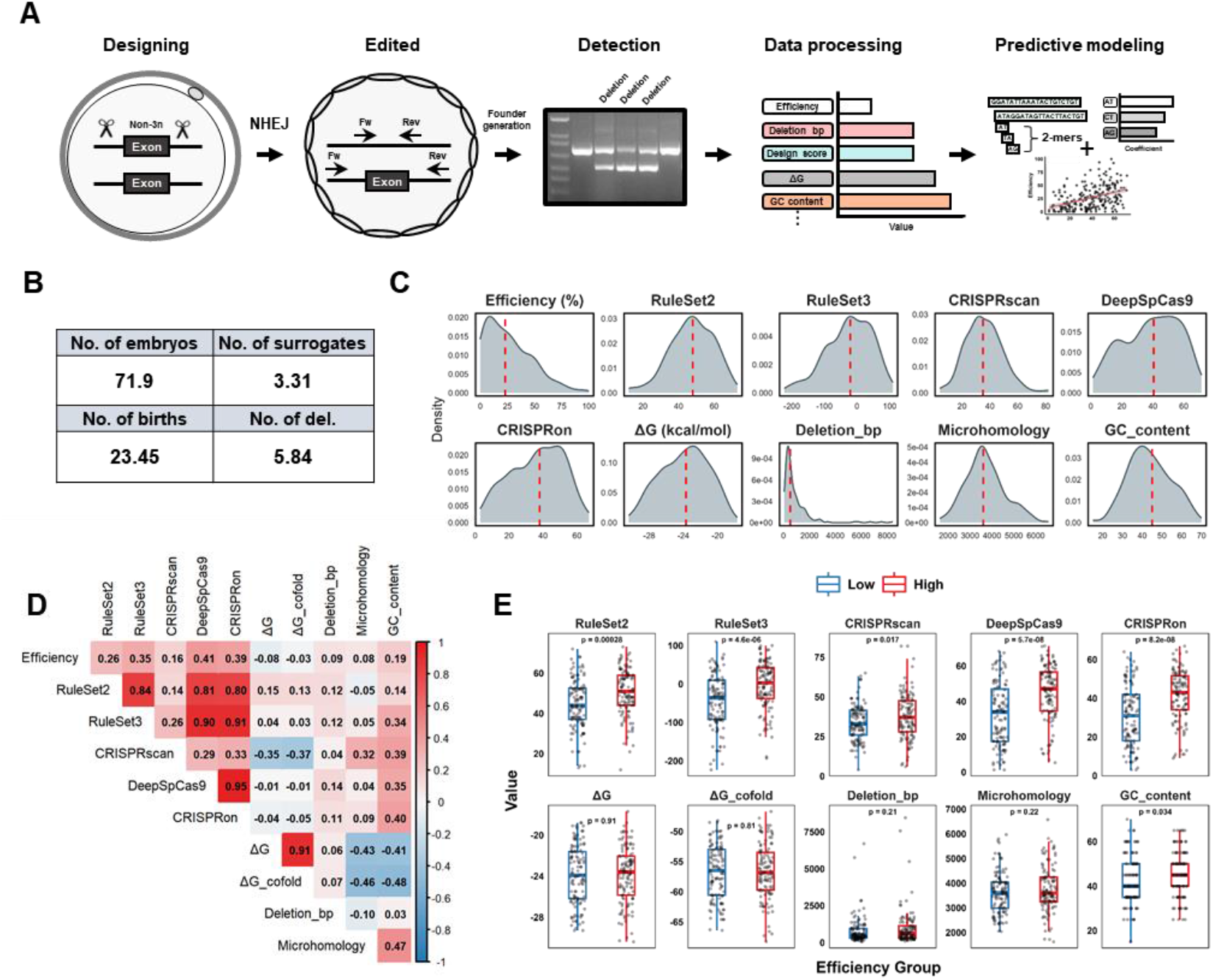
Overview of the workflow, data processing, and statistical analyses. **(A)** sgRNA pairs were designed to target exonic regions and induce indels via non-homologous end joining (NHEJ). Edited founders were screened and validated through gel electrophoresis. Quantified features were used for predictive modeling based on sgRNA sequence and design parameters. **(B)** Mouse generation information used for in vivo data. **(C)** Visualization of the distribution of data variables, including in vivo efficacy. The red line indicates the median. ΔG represents the Gibbs free energy change of the sgRNA. **(D)** Pearson’s correlation analysis of 10 features. ΔG cofold represents the Gibbs free energy change associated with the cofolding of two sgRNA molecules. **(E)** Comparative statistical analysis of 10 features. The Low and High groups were defined based on the median observed efficiency, and statistical significance was assessed using Welch’s t-test.

The medians of the 10 variables included in the analysis excluding strand orientation are as follows: Editing efficiency (23.03%), RuleSet2 (48), RuleSet3 (−20.5), CRISPRscan (35), DeepSpCas9 (40.53), CRISPRon (38), ΔG (23.7 kcal/mol), Deletion bp (518 bp), Microhomology (3604.55), and GC content (45%). In addition, all variables except for editing efficiency and deletion size exhibited a normal distribution pattern (**Figure 1C**). Among these variables, DeepSpCas9 (0.41) showed the highest Pearson correlation with editing efficiency. DeepSpCas9 also showed strong correlations (above 0.9) with RuleSet3 and CRISPRon (**Figure 1D**).

To further assess the significance of these associations, the data were divided into high-efficiency and low-efficiency groups based on the median editing efficiency, and Welch’s t-test was performed. As a result, DeepSpCas9, which had the highest correlation coefficient, showed the most statistically significant difference between the two groups (p = 5.7e−08). In addition, the following variables were found to be statistically significant based on p < 0.05: RuleSet2 (p = 2.8e-04), RuleSet3 (p = 4.6e−06), CRISPRscan (p = 0.017), CRISPRon (p = 8.2e−08), and GC content (p = 0.034) (**Figure 1E**). Regarding strand orientation, pairwise Wilcoxon rank-sum tests (Wilcoxon t-tests) indicated that there were no statistically significant differences across the three groups (**Supplementary Figure S1**).

### Predictive modeling

To reduce overfitting caused by design-related variables, the most statistically significant feature identified through correlation and association analyses—the DeepSpCas9 score—was selected as the sole input variable. Visualization of the relationship between DeepSpCas9 scores and editing efficiency revealed a weak linear association, with an R^2^ of 0.169 (**Figure 2A**).

**Figure 2.**
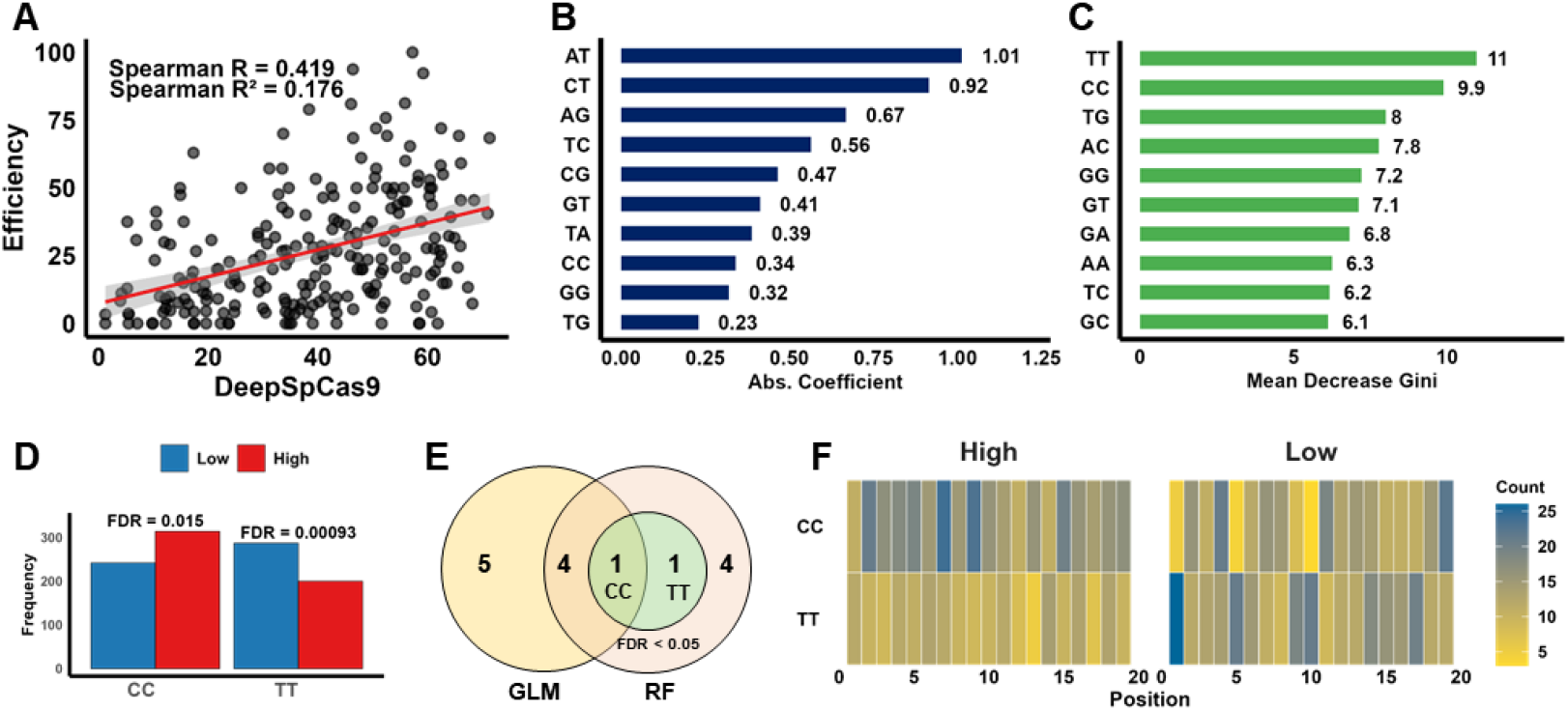
Profiling of selected design and K-mer-based features. **(A)** Scatter plot showing the distribution of the selected feature (SpDeepCas9 score) against observed editing efficiency, along with the Spearman correlation coefficient and R-squared value. **(B)** Visualization of the top 10 k-mers (k = 2) based on the absolute values of coefficients from the GLM model. **(C)** Visualization of the top 10 k-mers (k = 2) ranked by Mean Decrease in Gini from the Random Forest model. **(D)** Bar plots representing the frequencies of statistically significant 2-mers identified by GLM analysis, highlighting differential representation between high- and low-efficiency groups (FDR < 0.05, Fisher’s exact test with Benjamini–Hochberg correction). **(E)** Venn diagram showing the overlap of important k-mers identified by Generalized Linear Model (GLM), Random Forest (RF), and Fisher’s exact test (FDR < 0.05). **(F)** Heatmap displaying the positional distribution of statistically significant k-mers (FDR < 0.05) across low- and high-efficiency sgRNAs.

The GLM model trained using this variable (hereafter referred to as the Design GLM) achieved an AUC of 0.696, while the corresponding RF model (Design RF) showed an AUC of 0.65. For the k-mer-based models (k = 2), the GLM model (KMER GLM) achieved an AUC of 0.719, and the RF model (KMER RF) yielded an AUC of 0.654. Additionally, the Combined GLM (Design GLM + KMER GLM) and Combined RF (Design RF + KMER RF) models achieved AUCs of 0.759 and 0.684, respectively. In addition, two hybrid models, Hybrid 1 (Design GLM + KMER RF) and Hybrid 2 (Design RF + KMER GLM), showed AUCs of 0.713 and 0.726, respectively (**Supplementary Figure S2**).

The top three features in KMER GLM, ranked by absolute coefficient values, were AT, CT, and AG (**Figure 2B**), while in KMER RF, based on mean decrease in Gini, the top three were TT, CC, and TG (**Figure 2C**). The frequency of each k-mer was calculated in the high- and low-efficiency groups, and Fisher’s exact test followed by Benjamini–Hochberg correction was applied. Two k-mers, CC and TT, passed the FDR < 0.05 threshold and were considered significant (**Figure 2D**). A Venn diagram comparing the top 10 k-mer features selected by GLM and RF models showed that five features were commonly identified by both. CC (FDR < 0.05 in KMER GLM) appeared in both models, whereas TT (FDR < 0.001 in KMER GLM) was uniquely identified by the RF model (**Figure 2E**). To determine their positional distribution within crRNA sequences, a heatmap analysis was conducted. In the high efficiency group, CC was primarily enriched between positions 2 and 10, while TT was rare. In contrast, the low-efficiency group showed strong enrichment of TT at position 1 and relatively uniform distribution across the sequence, whereas CC displayed an opposing pattern (**Figure 2F**).

### Comparison of predictive models

To determine the optimal predictive model among the eight trained models (Design GLM, Design RF, KMER GLM, KMER RF, Combined GLM, Combined RF, Hybrid 1, and Hybrid 2), a visual comparison was conducted (**Figure 3A**), and to assess the similarity between the predicted probabilities of different models, both Pearson (**Figure 3B**) and Spearman (**Figure 3C**) correlation analyses were performed. As a result, the Combined GLM model consistently showed the highest values among the eight models. The top four model demonstrated the highest predictive performance. Results from repeated 10-fold cross-validation revealed the following mean AUCs (± standard deviation): Design GLM: 0.699 ± 0.12, Design RF: 0.659 ± 0.11, KMER GLM: 0.716 ± 0.11, KMER RF: 0.662 ± 0.11, Combined GLM: 0.759 ± 0.10, Combined RF: 0.687 ± 0.12, Hybrid 1: 0.715 ± 0.12, and Hybrid 2: 0.726 ± 0.11. Among the top four models with AUCs above 0.7 (Combined GLM, Hybrid 2, KMER GLM, and Hybrid 1), pairwise Welch’s t-tests showed statistically significant differences in performance (**Figure 3D**). However, DeLong’s test did not reveal statistically significant differences in AUC between models (p > 0.05) (**Supplementary Figure S3**).

**Figure 3.**
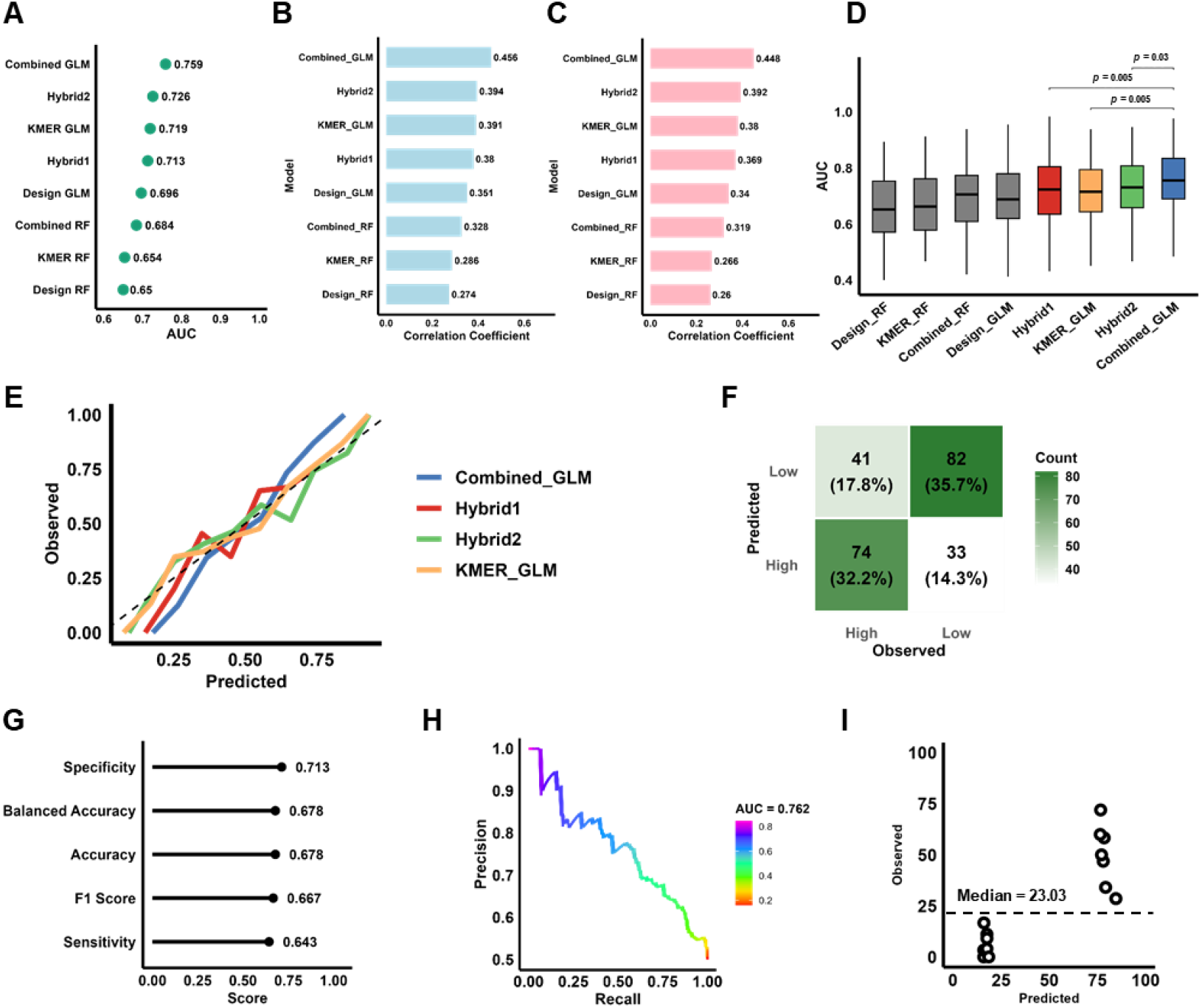
Model evaluation and selection. **(A)** Dot plot comparing the AUC values of eight predictive models, including Generalized Linear Model (GLM), Random Forest (RF) models trained on design features, k-mer features, combined features, and hybrid approaches. Area Under the Curve (AUC) values were calculated from Receiver Operating Characteristic (ROC) curves for each model to evaluate classification performance. **(B)** Pearson correlation coefficients between predicted values and true labels for each model, obtained via repeated 10-fold cross-validation. **(C)** Spearman correlation coefficients for each model based on repeated 10-fold cross-validation. **(D)** Box plot comparing AUC values from repeated 10-fold cross-validation across different models. Pairwise statistical comparisons were performed using Welch’s t-test, and p-values are displayed on the plot. **(E)** Calibration curves for different models, with predictions binned into 10 groups. **(F)** Confusion matrix for the Combined GLM model. Predictions were binarized at a threshold of 0.5 and compared to observed labels.**(G)** Summary of classification performance metrics for the Combined GLM model. Each dot represents a score based on the confusion matrix results. **(H)** Precision–Recall curve for the Combined GLM model. The curve shows the trade-off between precision and recall across various thresholds. The color gradient represents the threshold values, and the AUC is indicated in the legend. **(I)** Dot plot comparing predicted probabilities and observed efficiencies for the top and bottom 3% of sgRNAs based on Combined GLM predictions.

Calibration curve analysis further revealed that Combined GLM exhibited the best calibration among the models, indicating high reliability of predicted probabilities, while Hybrid 1 showed the poorest calibration performance (**Figure 3E**).

### Verification of the Combined GLM model

To further validate the Combined GLM model—which demonstrated the highest AUC and the best calibration performance—a confusion matrix analysis was conducted. In Combined GLM, the top three features with the largest absolute coefficient values were AT, CT, and AG, which were the same as those in the KMER GLM model (**Supplementary Figure S4A**). In the Combined RF model, the top three features based on the mean decrease in Gini were DeepSpCas9, TT, and CC, showing similar features to the KMER RF model (**Supplementary Figure S4B**). The model correctly classified approximately 68% of the total data (156 out of 230), indicating a reasonably reliable overall predictive accuracy (**Figure 3F**). Classification metrics showed a specificity of 0.678, sensitivity of 0.643, and both balanced accuracy and overall accuracy of 0.713. The F1 score, calculated as 2 × (precision × recall) / (precision + recall), was 0.667 (**Figure 3G**). In addition, the PR curve analysis yielded an AUC of 0.762, suggesting that the model maintained a favorable trade-off between precision and recall (**Figure 3H**). The Brier score was 0.2038 (**Supplementary Figure S5A**), and the result of the Hosmer–Lemeshow test (p = 0.2314) (**Supplementary Figure S5B**) indicates that the model fits the data adequately. Based on the Combined GLM predictions, analysis of the top 3% and bottom 3% of sgRNAs in terms of predicted probabilities and observed efficiencies showed that all seven efficiencies were classified according to the median value (23.03) (**Figure 3I**). The minimum predicted probability in the top 3% was 76.5%, while the maximum predicted probability in the bottom 3% was 18.6%.

## DISCUSSION

Effective sgRNA design is essential not only for achieving high gene editing efficiency but also for inducing a loss-of-function genotype. Considering that nonsense-mediated mRNA decay or truncated proteins typically arises from premature termination codon [29–31], the need for an in vivo–validated model for sgRNA design to exon deletion is underscored.

Previous studies have attempted to predict genome editing outcomes using in vitro or pre-implantation embryo data [32–34], but these approaches are limited by factors such as implantation failure, embryonic lethality, and mosaicism, which hinder their generalizability to whole organisms. To address these limitations, our study focused exclusively on viable postnatal individuals, establishing a more realistic in vivo benchmark. Even though the median editing efficiency of our data appears low at 23.03%, this reflects in vivo factors such as selective pressure, chromatin structure, and complexities in sgRNA processing [35–38].

Notably, none of the five intrinsic parameters of dual-sgRNAs predicted in vivo deletion efficiency, highlighting the limitations of conventional design features in biological contexts. Although our dataset was constrained by deletion sizes (50 bp–8.5 kb) and a non-normal distribution, recent evidence from an independent study supports the robustness of our findings [39].

Among eight models based on GLM and RF trained with 230 in vivo data, the final selected model—a Combined GLM— incorporated the DeepSpCas9 score (Pearson’s R = 0.41) and k-mer (k = 2) features. While these variables showed limited predictive power on their own, their combination achieved the best overall performance.

Interestingly, despite the TT and CC dinucleotides identified as significant motifs in the KMER GLM (**Figure 2D**) were not among the top features in terms of absolute coefficient values within the same model (**Figure 2B**), they emerged as some of the most important features in the RF model using the same input features (**Figure 2C**). This pattern was also maintained in both the Combined GLM and RF models (**Supplementary Figure S4A and S4B**). This discrepancy illustrates the methodological distinction between GLM, which focuses on linear and additive effects, and RF, which captures nonlinear interactions and higher-order dependencies.

In addition, k-mer positional distribution in sgRNAs showed that TT was enriched in sgRNAs with low editing efficiency. A plausible explanation is premature transcription termination during in vitro synthesis by T7 RNA polymerase, particularly when TT motifs are located near the 5′ end of the guide sequence [40].

Conversely, CC was enriched in high efficiency sgRNAs and may mark loci with greater chromatin accessibility. G-rich and CpG-rich sequences given strand symmetry often correlate with open chromatin features such as DNase I hypersensitivity and active histone modifications, which facilitate Cas9–RNP binding and cleavage [41]. While our study did not directly assess chromatin state, these results align with prior findings and suggest that short motifs like CC/GG may serve as proxies for epigenomic accessibility in vivo.

Although AUC of Combined GLM did not exceed 0.9, confidence interval analysis according to [42] showed that, under balanced binary classification (n_1_ = n_0_ = 115), the 95% confidence interval AUC value is approximately between 0.697 and 0.821. Therefore, AUC of 0.759 for the model’s performance is deemed statistically valid for this data size.

In conclusion, this study proposed that the combination of sequence-level features, and previous design scores enables more accurate prediction of CRISPR/Cas9 knockout efficiency in mammalian systems. These results challenge the assumption that design scores alone are sufficient and underscore the importance of modeling biologically relevant variables. Ultimately, the framework presented here may serve as a foundation for next-generation in vivo sgRNA design tools, with direct applications in animal model generation, gene therapy, and precision genome engineering.

## AUTHOR CONTRIBUTIONS

S-Y.L. supervised the project. S-Y.L. conceived the study (conceptualization, investigation, formal analysis). S-Y.L. performed bioinformatics analysis and modeling. S-Y.L. wrote the manuscript (data curation, methodology, visualization, writing original draft and editing). S-Y.L. performed sgRNA design and in silico analysis. S-Y.L., S.M., S.D.J., H.K., B.J., S-H.H., E.J., J.L., Y-K.L., and D.L. performed the in vitro and in vivo experiments. S-Y.L. reviewed and revised the manuscript. All authors have read and agreed to the published version of the manuscript.

## DATA AVAILABILITY

The final selected prediction model Combined GLM (CRISPRsy) is available at: https://sungyeonlee0711.shinyapps.io/dsgRNA_predictor/

## FIGURE LEGEND

**Supplementary Figure S1. Comparison of editing efficiency by sgRNA strand orientation**.Efficiency distributions are shown for Convergent (n = 59), Divergent (n = 61), and Same (n = 110) sgRNA pairs. Statistical significance was assessed using the Wilcoxon rank sum test with Benjamini– Hochberg correction; adjusted p-values are shown above the relevant comparisons.

**Supplementary Figure S2. ROC curves of the eight generated predictive models**.

**Supplementary Figure S3. Comparison of AUC values between models using the DeLong test**. Each dot represents a pairwise comparison, with –log_10_ (p-value) on the x-axis. The dashed line indicates the significance threshold (p = 0.05). Raw p-values are shown next to each point.

**Supplementary Figure S4. Top 10 features ranked by average importance from Generalized Linear Model (GLM) and Random Forest (RF) models. (A)** Top 10 features based on GLM importance. Features were selected by averaging the absolute coefficients from two separate GLM models trained on design and k-mer features, respectively. **(B)** Top 10 features based on RF importance. Mean Decrease in Gini values were averaged across models trained on design and k mer features to determine overall feature relevance.

**Supplementary Figure S5. Calibration evaluation of the Combined GLM model. (A)** Brier Score for the Combined GLM model. **(B)** Hosmer–Lemeshow calibration plot for the Combined GLM model (p = 0.23).

