## Supplementary figures and images for "Empirical optimization of dual-sgRNA design for in vivo CRISPR/Cas9-mediated exon deletion in mice"

### Supplementary Figure S1

#### Supplementary Figure S1

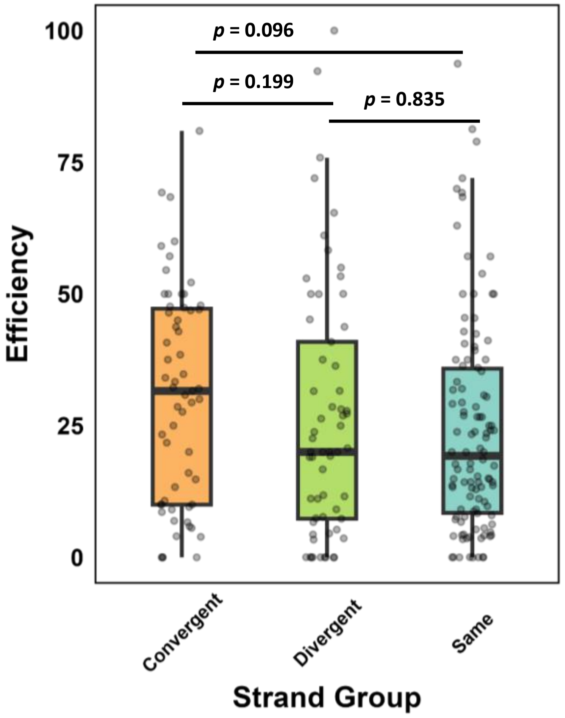

### Supplementary Figure S2

#### Supplementary Figure S2

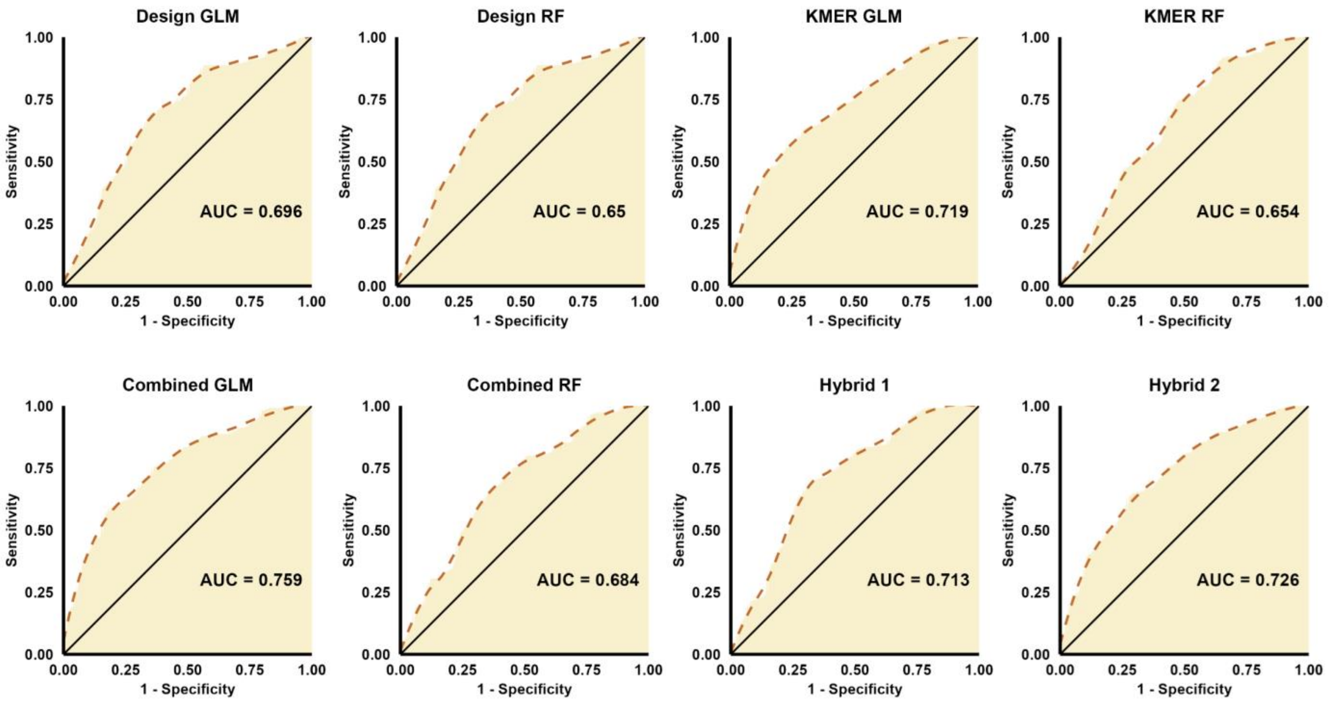

### Supplementary Figure S3

#### Supplementary Figure S3

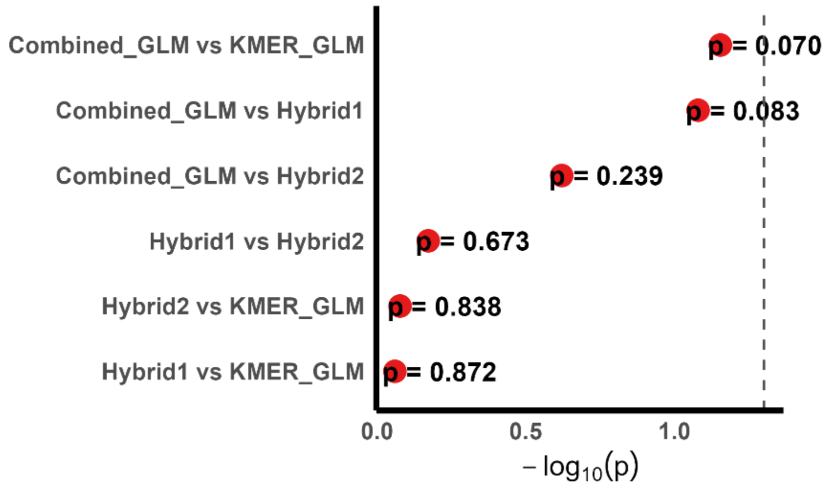

### Supplementary Figure S4

#### Supplementary Figure S4

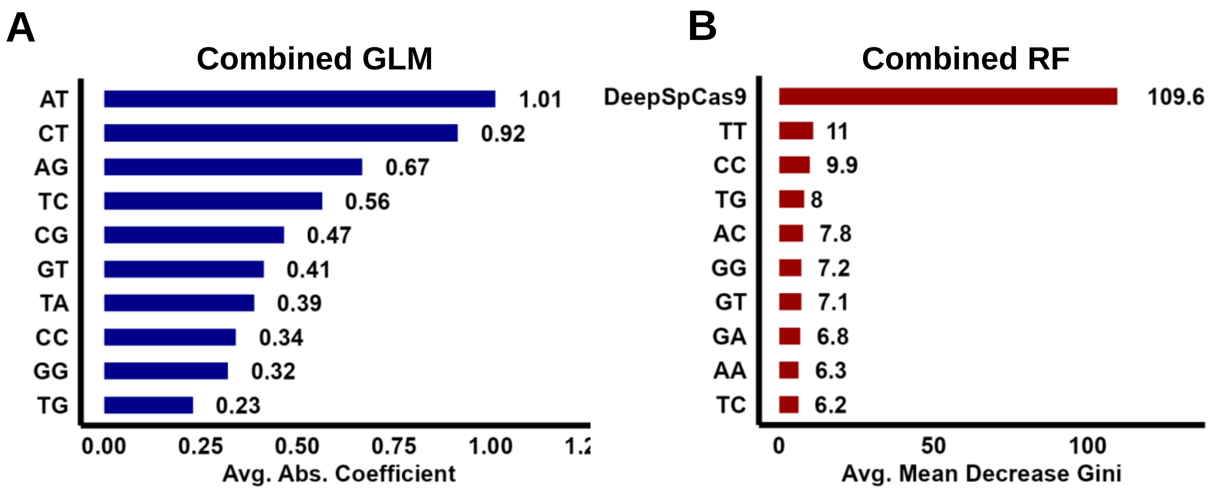

### Supplementary Figure S5

##### Supplementary Figure S5

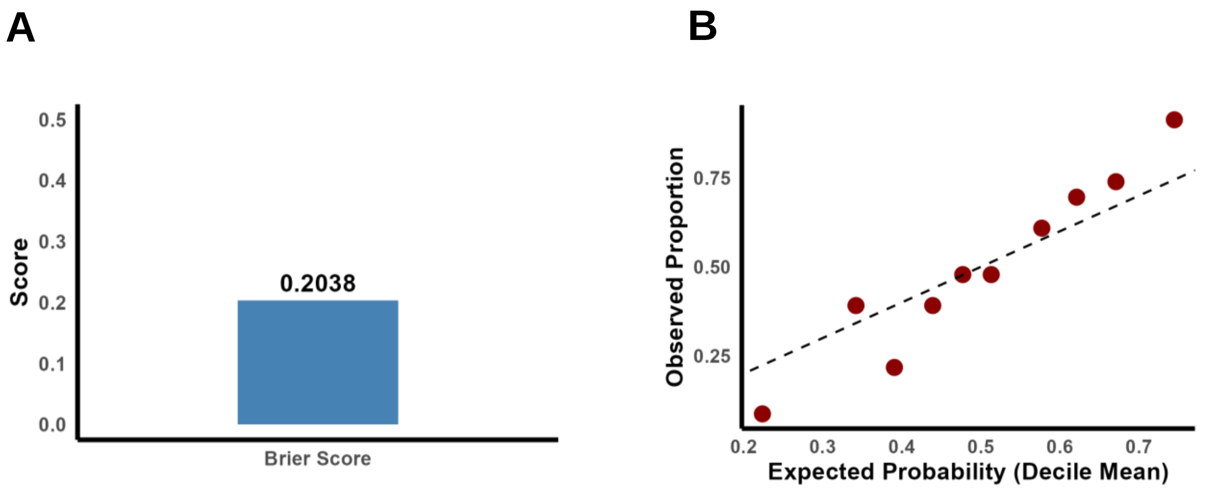
